# Autoinhibition and conformational activation regulate cell-wall anchoring of the bacterial type IV pilus machinery

**DOI:** 10.64898/2026.11.18.767550

**Authors:** Hilario Domínguez-Cerván, Laura Barrientos-Moreno, Juan Fernández-Carrillo, Carmen González-Molina, Inés G. Muñoz, Cayo Ramos, Inmaculada Pérez-Dorado, Luis Rodríguez-Moreno

## Abstract

Type IV pili are widespread bacterial nanomachines that mediate adhesion, motility and host interactions, yet the molecular mechanism coordinating assembly of their outer-membrane channel with attachment to the bacterial cell wall has remained unknown. Here, we show that the conserved PilQ-associated protein TsaP functions as a conformationally regulated envelope adaptor linking secretin assembly to cell-wall anchoring. A crystal structure of full-length TsaP, integrated with structural, biophysical and functional analyses, reveals a compact autoinhibited conformation in which the N-terminal LysM domain is sequestered against the protein core. Comparative structural analyses support a mechanism in which PilQ association remodels TsaP, exposing the LysM domain for peptidoglycan binding. Together, our findings support a mechanistic model in which PilQ-dependent conformational activation of TsaP couples secretin assembly to cell-wall anchoring, providing a framework for understanding the regulated integration of type IV pilus machineries within the bacterial envelope.

## Introduction

Type IV pili (T4P) are widespread bacterial surface appendages that mediate a broad range of functions including adhesion, twitching motility, biofilm formation, surface sensing, DNA uptake and host colonization (Denise *et al*, 2019; Craig *et al*, 2019; Ellison *et al*, 2022; Gaines *et al*, 2024). As members of the type IV filament (TFF) superfamily, T4P are assembled by complex envelope-spanning nanomachines that have diversified throughout Bacteria and Archaea while retaining a conserved architectural framework (Craig *et al*, 2004; Denise *et al*, 2019; Gaines *et al*, 2022; Liu *et al*, 2024). In Gram-negative bacteria, T4P biogenesis requires the coordinated action of inner-membrane assembly ATPases, periplasmic platform proteins, and the outer-membrane secretin PilQ, which forms the conduit through which the pilus is extruded (Gold & Kudryashev, 2016; Hospenthal *et al*, 2017). The assembly and long-term stability of these large supramolecular complexes rely on accessory proteins that integrate the machinery within the cell envelope and ensure its mechanical coupling to the surrounding cell wall (Siewering *et al*, 2014; Mccallum *et al*, 2019; Tassinari *et al*, 2023).

Among these accessory factors, TsaP (T4P secretin-associated protein) has emerged as a conserved component associated with the secretin complex in several T4P systems (Siewering *et al*, 2014). Structural and genetic studies indicate that TsaP forms oligomeric assemblies around PilQ and contributes to the architecture and stability of the outer-membrane/periplasmic region of the pilus machinery. Cryo-electron microscopy (cryo-EM) analyses of PilQ-containing complexes from different bacterial species have revealed that TsaP adopts ring-like arrangements surrounding the secretin barrel, suggesting an important role in coordinating secretin organization within the envelope (Chang *et al*, 2016; McCallum *et al*, 2021). In addition, the presence of an N-terminal LysM domain, a widespread peptidoglycan (PG)-binding module, has led to the proposal that TsaP contributes to anchoring the T4P machinery to the PG layer (Siewering *et al*, 2014).

Despite increasing structural information on PilQ-TsaP assemblies, the molecular basis of this proposed PG-anchoring function of TsaP remains poorly understood. Available cryo-EM reconstructions have primarily captured TsaP within secretin-associated complexes and provide only partial information on the organization of the full-length protein (Guo *et al*, 2024; McCallum *et al*, 2021). Consequently, it remains unclear how the LysM domain is positioned within TsaP, how PG recognition is regulated, and whether secretin binding influences the functional state of the protein. More broadly, the mechanisms that coordinate secretin assembly with cell-wall engagement during T4P biogenesis remain largely unknown.

*Pseudomonas savastanoi* pv. savastanoi (Psv), the causal agent of olive knot disease, encodes a LysM-containing protein, originally designated LysM4, that we previously identified as the TsaP orthologue required for bacterial virulence and host colonization (Domínguez-Cerván *et al*, 2025). To reflect its evolutionary and functional conservation, we hereafter refer to this protein as PsvTsaP. The close evolutionary and functional relationship between PsvTsaP and TsaP proteins from other *Pseudomonas* species makes this system particularly suitable for investigating the molecular mechanisms that coordinate secretin assembly with PG engagement.

Here, we define the architecture of full-length PsvTsaP and investigate the molecular mechanism underlying its conformational regulation using X-ray crystallography, small-angle X-ray scattering, biophysical and glycan-binding analyses, structure-guided computational modelling and bacterial genetics. Our results reveal that TsaP adopts a compact autoinhibited conformation in which the LysM domain is sequestered against the protein core, generating a PG-locked state that restricts premature cell-wall engagement. Structural and functional analyses further support a model in which association with PilQ promotes repositioning of the LysM domain and exposure of its glycan-binding surface. Together, these findings provide a structural framework for understanding how secretin-associated envelope adaptors are spatially and conformationally regulated during T4P assembly. More broadly, they support the view that TsaP-like proteins are not passive tethers but regulated architectural elements that coordinate the ordered assembly and envelope integration of T4P machineries.

## METHODS

### Bacterial strains and growth conditions

The bacterial strains and plasmids used in this study are listed in table S1. *Escherichia coli* was routinely grown at 37°C in lysogeny (LB) medium (BERTANI, 1951) without glucose and containing 0.5 %(w/v) NaCl. Psv NCPPB 3335 and its derivative mutants were routinely grown in Kinǵs B (KB) medium (King *et al*, 1954) at 28°C. When required, kanamycin (Km) was added at a final concentration of 50 µg/mL.

### Construct design and protein expression

Constructs encoding the mature full-length PsvTsaP (residues 21-341; UniProt: A0ABC8B5Y8) and its sole LysM domain (PsvLysM, residues 19-89) were cloned into a pET28a, respectively (details in table S1). The resulting recombinant plasmids (pET28a::PsvTsaP and pET28a::PsvLysM) were used to transform *E. coli* BL21 GOLD (DE3) pLysS cells for protein expression in 1 L cultures of LB medium supplemented with Km. Cells were grown at 37°C until an optical density at 600 nm (OD_600_) of 0.5-0.8, followed by induction of protein expression with 0.2 mM addition of isopropyl β-D-1-thiogalactopyranoside (IPTG) and incubation at 18°C for 24 h. Cells were harvested by centrifugation at 5251 *g* for 30 min at 4°C.

### Purification of full-length PsvTsaP under denaturing conditions

Purification of His-tagged PsvTsaP under denaturing conditions was performed following a modified version of the protocol described by Mammeri et al., 2019. Cell pellets were resuspended in denaturing buffer (50 mM Tris-HCl, 500 mM NaCl, 6 M guanidine hydrochloride) and incubated at 60°C until complete solubilization. The lysate was then clarified by centrifugation at 100,000 *g* for 1 h at 16°C using an R30AT Himac CR30NX ultracentrifuge. Solubilized inclusion bodies were subjected to immobilized metal affinity chromatography (IMAC) using a 5 mL HisTrap HP column (Cytiva, Marlborough, Massachusetts, USA) using an ÄKTA Go FPLC system previously equilibrated with binding buffer (50 mM Tris-HCl, 500 mM NaCl, 20 mM imidazole, 8 M urea, pH 8.0). Bound proteins were eluted using a linear gradient of elution buffer (50 mM Tris-HCl, 500 mM NaCl, 500 mM imidazole, 8 M urea, pH 8.0). After affinity purification, the eluted protein was dialyzed using Spectra/Por 4 dialysis membrane standard RC tubing (Spectrumlabs, San Francisco, California, USA) to exchange the elution buffer for size exclusion chromatography (SEC) buffer (20 mM Tris-HCl pH 8.0, 150 mM NaCl). Pure fractions containing the PsvTsaP were concentrated using Amicon Ultra centrifugal filters (Merck Millipore, Dublin, Ireland), followed by SEC on a HiLoad 16/600 Superdex 200 pg column (Cytiva, Marlborough, Massachusetts, USA) equilibrated with SEC buffer.

### Purification of His-tagged PsvLysM domain

Cell pellets from IPTG-induced cultures expressing PsvLysM were resuspended in lysis buffer (50 mM Tris-HCl pH 8.0, 500 mM NaCl, 10 mM imidazole, 10 mM DNase, 1 mM MnCl_2_, 10 mM MgCl_2_) followed by a 15 min incubation at room temperature. Cells were lysed by sonication on ice using an Ultrasonic processor (Sigma-Aldrich, Saint Louis, Missouri, USA) on ice (15 min total, 15 s pulses at 40 % power). The lysate was then clarified by centrifugation at 100,000 *g* for 1 h at 4°C using an R30AT Himac CR30NX ultracentrifuge. The clarified lysate was subjected to IMAC using a 5 mL HisTrap HP column (Cytiva, Marlborough, Massachusetts, USA) using an ÄKTA Go FPLC system. The column was washed with five column volumes of binding buffer (50 mM Tris-HCl pH 8.0, 500 mM NaCl, 10 mM imidazole), followed by protein elution with a linear gradient of elution buffer (50 mM Tris-HCl pH 8.0, 500 mM NaCl, 500 mM imidazole). Fractions containing pure His-tagged PsvLysM domain were concentrated using Amicon Ultra centrifugal filters (Merck Millipore, Dublin, Ireland) and further purified by SEC on a HiLoad 16/600 Superdex 200 pg column (Cytiva, Marlborough, Massachusetts, USA) equilibrated with SEC buffer (20 mM Tris-HCl pH 8.0, 150 mM NaCl). For His-tag removal, the protein was incubated overnight with HRV 3C protease at a protease:target protein ratio of 1:100 (w/w) in cleavage buffer (20 mM Tris-HCl pH 8.0, 150 mM NaCl) at 4°C under gentle agitation. The untagged PsvLysM was subjected to SEC on a HiLoad 16/600 Superdex 200 pg column (Cytiva, Marlborough, Massachusetts, USA) equilibrated with SEC buffer. In all cases, pure protein fractions from the SEC were pooled, concentrated, flash frozen in liquid nitrogen, and stored at −80°C.

### Differential scanning fluorimetry (nanoDSF) assays

NanoDSF assays were performed to determine the inflection temperature (*T*_i_) associated with the thermal unfolding transition of PsvTsaP and the sole PsvLysM domain in the presence or absence of hexa-*N*-acetylglucosamine (NAG_6_). Measurements were carried out using a Tycho NT.6 instrument (NanoTemper Technologies GmbH, Munich, Germany) equipped with standard capillaries provided by the manufacturer. Each reaction was prepared in a total volume of 10 μL in DSF buffer (20 mM HEPES, pH 8.0, 150 mM NaCl) containing 5 μM purified protein. Samples supplemented with ligand contained 1 mM NAG_6_. Each condition was analyzed in triplicate, and mean *T*_i_ values and standard deviations were calculated from three independent measurements. Graphs were generated using GraphPad Prism 8.

### Crystallization, X-ray data collection and structure determination

Crystallization experiments were performed by sitting-drop vapor diffusion method at 18°C using 96-well MRC plates (Hampton Research, Aliso Viejo, California, USA) and an Oryx8 crystallization robot (Douglas Instruments, Berkshire, UK). Crystals of PsvTsaP grew in 300 nL drops formed by mixing 150 nL of protein solution at 6.25 mg/mL with 150 nL of precipitant solution (25 mM HEPES pH 7.0 and 0.8 M Na^+^/K^+^ tartrate). Crystals were flash frozen in liquid nitrogen without cryoprotection and subsequently used to measure a complete X-ray data set at 100 K in the beamline ID23 at ESRF (Grenoble, France). Diffraction data were integrated, scaled, and reduced using XDS (Kabsch, 2010) and AIMLESS (Evans & Murshudov, 2013). Data collection statistics are summarized in table S2. The structure of PsvTsaP was solved by the molecular replacement method with PHASER (McCoy *et al*, 2007) and using an AlphaFold2-predicted (Jumper *et al*, 2021) model of PsvTsaP as the search model. The initial model was refined through iterative cycles of manual building with COOT (Emsley *et al*, 2010) and refinement using PHENIX (Adams *et al*, 2010). Figures were prepared using PyMOL (Schrödinger, Cambridge, Massachussetts, USA). The atomic coordinates and structure factors have been deposited in the Protein Data Bank (PDB) under the entry 31GV.

### Small-angle X-ray scattering (SAXS)

SAXS experiments were carried out at the B21 beamline of Diamond Light Source (Harwell Campus, UK) (Cowieson *et al*, 2020). A total of 45 μL of His-tagged PsvTsaP protein at 6.5 mg/mL were injected onto an SRT-C SEC-300 (Sepax, Newark, Delaware, USA) column equilibrated in buffer containing 20 mM Tris-HCl pH 8.0, 1 % (v/v) glycerol, and 150 mM NaCl, and run on an Agilent 1200 HPLC system at 18°C. Continuously eluting fractions were exposed for 3 s within 10 s acquisition cycles using an X-ray wavelength of 1 Å and a sample-detector distance of 3.7 m on an Eiger 4M detector. The collected data spanned a momentum transfer range of 0.0032 < q < 0.34 Å^-1^. Frames acquired immediately before sample elution were used for buffer subtraction. Data analysis, including subtraction, scaling, merging, and assessment of radiation damage, was performed using the scÅtter software package (www.bioisis.net). The radius of gyration (Rg) was determined by Guinier analysis under the assumption q < 1.3/Rg. The particle distance distribution and maximum dimension (Dmax) were obtained with GNOM, and *ab initio* shape reconstructions were generated with DAMMIF/DAMMIN, all from the ATSAS package (Manalastas-Cantos *et al*, 2021). The protein molecular mass was also estimated using GNOM. Atomic models based on the X-ray structure determined in this study were manually fitted into the SAXS-derived envelope in COOT (Emsley *et al*, 2010), and the agreement between the experimental scattering profile and the fitted model was evaluated with FoXS (Schneidman-Duhovny *et al*, 2016). Data collection statistics are summarized in table S3.

### Sequence analysis

Sequence alignments were performed with Clustal Omega (Madeira *et al*, 2024) using default parameters. The sequence of PsvTsaP (UniProt: A0ABC8B5Y8) was aligned with TsaP homologues from *Pseudomonas aeruginosa* (PaTsaP; UniProt: Q9I7A7), *Pseudomonas oleovorans* (PlvTsaP; UniProt: A0A2T5PN03), *Vibrio cholerae* (VchTsaP; UniProt: Q9KVU2), *Legionella pneumophila* (LpnTsaP; UniProt: Q5ZSC3), *Acinetobacter baumannii* (AbmTsaP; UniProt: A0A7U4DF81), *Myxococcus xanthus* (MxnTsaP; UniProt: Q1D813) and *Neisseria gonorrhoeae* (NgnTsaP; UniProt: A0AB74EQ34). The sequence of the PsvLysM domain was aligned with the LysM regions of the TsaP homologues listed above and with representative LysM-containing proteins of known structures from bacterial and fungal species, as detailed below in structural comparisons and homology searches.

The sequence of PsvPilQ (UniProt: A0ABC8BJ56) was aligned with representative PilQ homologous from diverse bacteria including those from *P. aeruginosa* (PaPilQ; UniProt: P34750), *Azotobacter beijerinckii* (AbjPilQ; UniProt: A0A1I3YC62), *P. oleovorans* (PlvPilQ; UniProt: A0A653B6Z8), *V. cholerae* (VchPilQ; UniProt: A0A655VTG1), *L. pneumophila* (LpnPilQ; UniProt: A0A3A6W2J6), *A. baumannii* (AbmPilQ; UniProt: A0A6F8TC36), *M. xanthus* (MxnPilQ; UniProt: Q9ZFG1), *N. gonorrhoeae* (NgnPilQ, UniProt: P35819).

### Structural comparisons and homology searches

Structural homology searches for PsvTsaP were performed using Foldseek (van Kempen *et al*, 2024). Structural comparisons of the PsvLysM domain with representative LysM-containing proteins were carried out using experimentally determined structures from the PDB, including Ecp6 from *Cladosporium fulvum* (CflEcp6; PDB: 4B8V), and NlpC/P60 from *Thermus thermophilus* (TthNlpC/P60; PDB: 4UZ3). Structural superpositions were performed using the RCSB PDB structure comparison tool and PyMOL (Schrödinger, Cambridge, Massachussetts, USA).

### Computational modelling of three-dimensional structures

The three-dimensional structure of computational models was predicted using AlphaFold 3 (AF3) (Abramson *et al*, 2024). Firstly, the structure of TsaP from *N. gonorrhoeae* (NgnTsaP, residues 50-415; UniProt: A0AB74EQ34) was modelled using the AlphaFold Server (Google DeepMind/Isomorphic Labs) using default parameters. Secondly, a representative complex of a PsvTsaP:PilQ assembly, here referred to as PsvTsaP:PilQ complex, consisting of 4 copies of PsvPilQ (residues 444-701) bound to one copy of PsvTsaP (residues 112-341, corresponding to CI and CII domains), was modelled using the AlphaFold Server with default parameters. Thirdly, the structure of the PsvLysM domain bound to the PG analogue *N*-acetylmuramyl β(1→4) *N*-acetylglucosamyl β(1→4) *N*-acetylmuramyl β(1→4) *N*-acetylglucosaminide ([NAM-NAG]_2_), referred to as PsvLysM:[NAM-NAG]_2_ complex, was modelled using a local AF3 installation. To do so, two input files were fed to AF3 consisting of the PsvLysM sequence (residues 19-89), and a JSON (alphafold3 dialect, version 1) file including the definitions of the [NAM-NAG]_2_ molecule. The latter includes: i) the alternating *N*-acetylmuramic acid and *N*-acetylglucosamine of the ligand, specified using RCSB PDB CCD codes; and ii) three β(1→4) glycosidic bonds present in the ligand that define the connection between C1 and O4 atoms of acceptor/donor saccharide rings, via the *bondedAtomPairs* field of the JSON file, to preserve monosaccharide and linkage stereochemistry(Huang *et al*, 2025). A single seed (seed = 1) was used. The reliability of PsvTsaP:PilQ and PsvLysM:[NAM-NAG]_2_ complexes was evaluated based on the top-ranked model retained and confidence metrics (ipTM, pTM, PAE and pLDDT) extracted from the JSON output (Abramson *et al*, 2024).

## RESULTS

### PsvTsaP adopts a compact multidomain architecture associated with type IV pilus assembly

PsvTsaP has been implicated in host-tissue colonization by Psv and predicted to associate with the T4P machinery (Domínguez-Cerván *et al*, 2025). Consistent with this proposed role, PsvTsaP displays high sequence similarity to the TsaP from *P. aeruginosa* and other TsaP homologues (Appendix Fig. S1A). To gain structural insight into PsvTsaP function, we determined a crystal structure of full-length PsvTsaP at 2.74 Å resolution (Fig. 1B and Appendix Fig S1B). The structure shows that PsvTsaP arranges as a multi-domain protein comprising an N-terminal LysM domain followed by two consecutive β-barrel domains referred to as CI and CII. LysM and CI/II domains are connected by long inter-domain linker regions referred to as Linker 1 (L1), which includes two short α-helices, and Linker 2 (L2) that is partially flexible and not visible in the structure (Fig. 1A, B).

**Figure 1.**
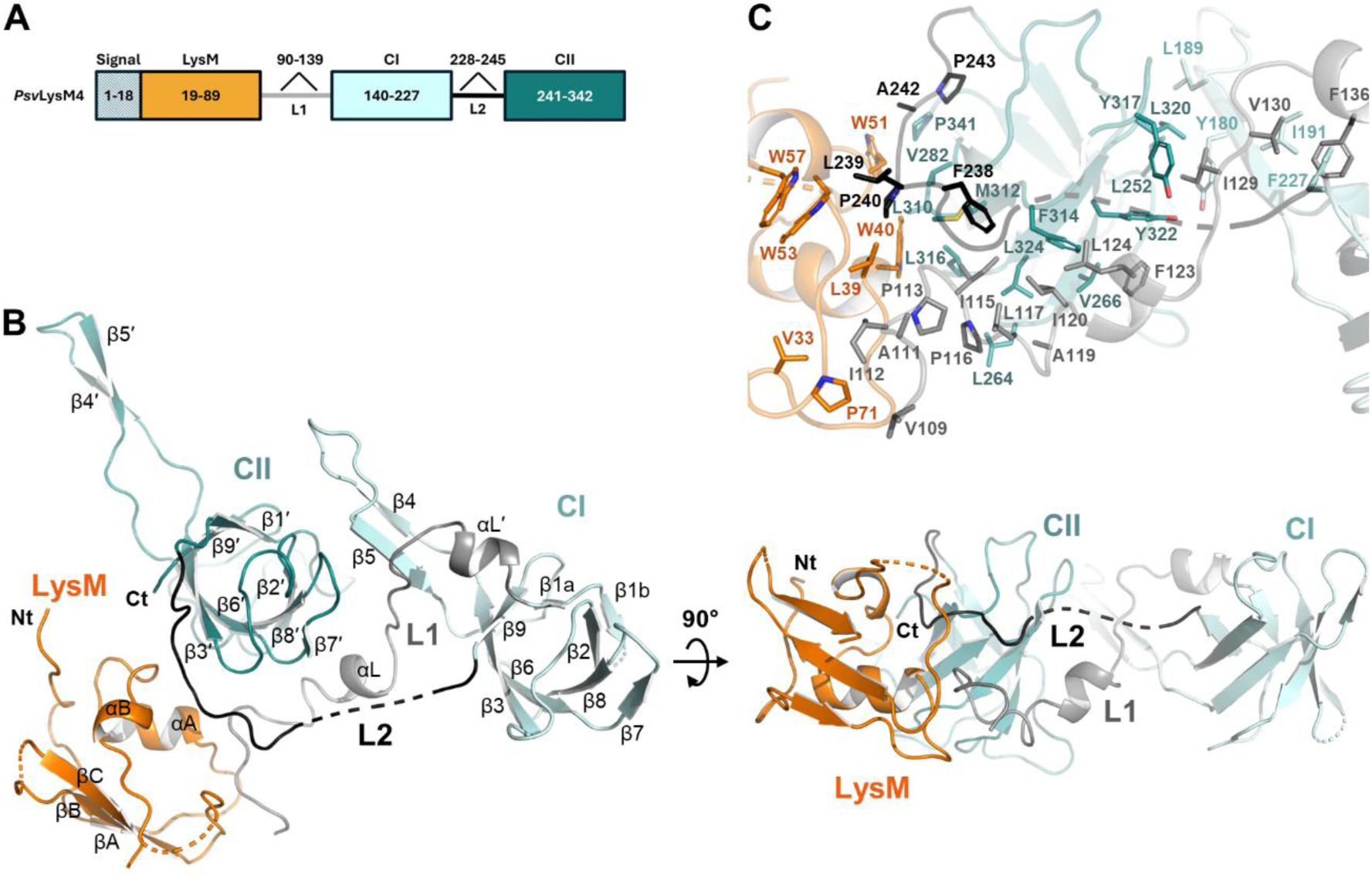
PsvTsaP crystal structure and effect on Psv surface appendages. (**A**) Domain architecture of the mature full-length PsvTsaP, showing the signal peptide (grey), the N-terminal LysM domain (in orange), the L1 and L2 linkers (in grey and black, respectively), and the two C-terminal β-barrel domains, CI and CII (in light and dark teal, respectively). Residue boundaries are indicated. (**B**) Orthogonal views of full-length PsvTsaP crystal structure represented in cartoons and coloured using the same code as in A. (**C**) Close-up view of the LysM-CI/CII interface. Residues contributing to the inter-domain contacts are shown as sticks and labelled as in A and B.

The LysM domain exhibits a conserved βααβ-fold characteristic of this class of motifs, although the antiparallel two-stranded β-sheet is extended by a third β-strand (βC) at the C-terminus (Fig. 1B, and fig. S1A). CI and CII are two consecutive small domains exhibiting a conserved six-stranded antiparallel β-barrel core (RMSD = 2.07 Å for 56 Cα atoms), which are capped by four loops, two at each side (Fig. 1B and Appendix Fig. S1A, D). Both CI and CII domains are also decorated with a long extension accommodating a β-hairpin formed by β4-β5 and β4’-β5’ strands, respectively. Rather than behaving as independent modules, the three globular domains are tightly fastened into a compact architecture stabilized by extensive interdomain and linker-mediated interactions (Fig. 1C). These include direct inter-domain LysM-CII and CI-CII contacts where the CI β-hairpin fastens CII β-barrel adopting a head-to-tail arrangement. In addition, linker-mediated interactions further bridge the three globular domains through an extensive network of contacts established across L1 and L2 connectors. The latter is primarily sustained by hydrophobic interactions involving residues exhibiting high or partial conservation across homologue TsaP sequences inspected; and one salt-bridge, moderately conserved in sequence (Fig. 1C and Appendix Fig. S1A, C).

The high degree of sequence homology between PsvTsaP and PaTsaP extends to a high conservation of their three-dimensional structure (Appendix Fig. S1E). This is evidenced by the superimposition of PsvTsaP with the partial structure of PaTsaP within secretin-associated complexes determined by cryo-EM (PDB: 6VE2; McCallum et al., 2021) (RMSD = 2.974 Å for 165 Cα atoms) (Appendix Fig. S1E), which supports PsvTsaP as a TsaP-like protein suited for secretin interaction during T4P complex assembly in Psv.

### PsvTsaP adopts a PG-locked conformation regulating protein location at the bacterial cell-wall

The crystal structure of PsvTsaP shows an unexpected arrangement of the LysM domain, which interacts through its putative PG-binding site with residues 111-114 of the linker L1 (Fig. 2A). These LysM:L1 interactions result in a non-productive conformation for cell-wall binding, here referred to as a PG-locked conformation. This arrangement is stabilized by interactions involving conserved residues, including hydrogen bonds formed between the main-chain atoms of I69 and I112, and of L39 and P113, together with hydrophobic interactions between the side chains of V33, A35, L39 and P71 (in PsvLysM), and V109 and I112 (in L1). This PG-locked conformation is further sustained by numerous inter-domain interactions connecting the PG-binding site of PsvLysM with CII and L1/L2 regions (Fig. 2A). This PG-locked arrangement contrasts with the cryo-EM T4P structures from *P. aeruginosa* (PaT4P; PDB: 6VE2) and *M. xanthus* (MxnT4P; PDB: 3JC9), where the LysM domains are either not visible or located away from CI/II domains, respectively. As suggested by these studies, L1 and LysM appear to behave as flexible regions that may disengage from the CI/CII core, thereby repositioning the LysM domain into a conformation compatible with PG binding.

**Figure. 2.**
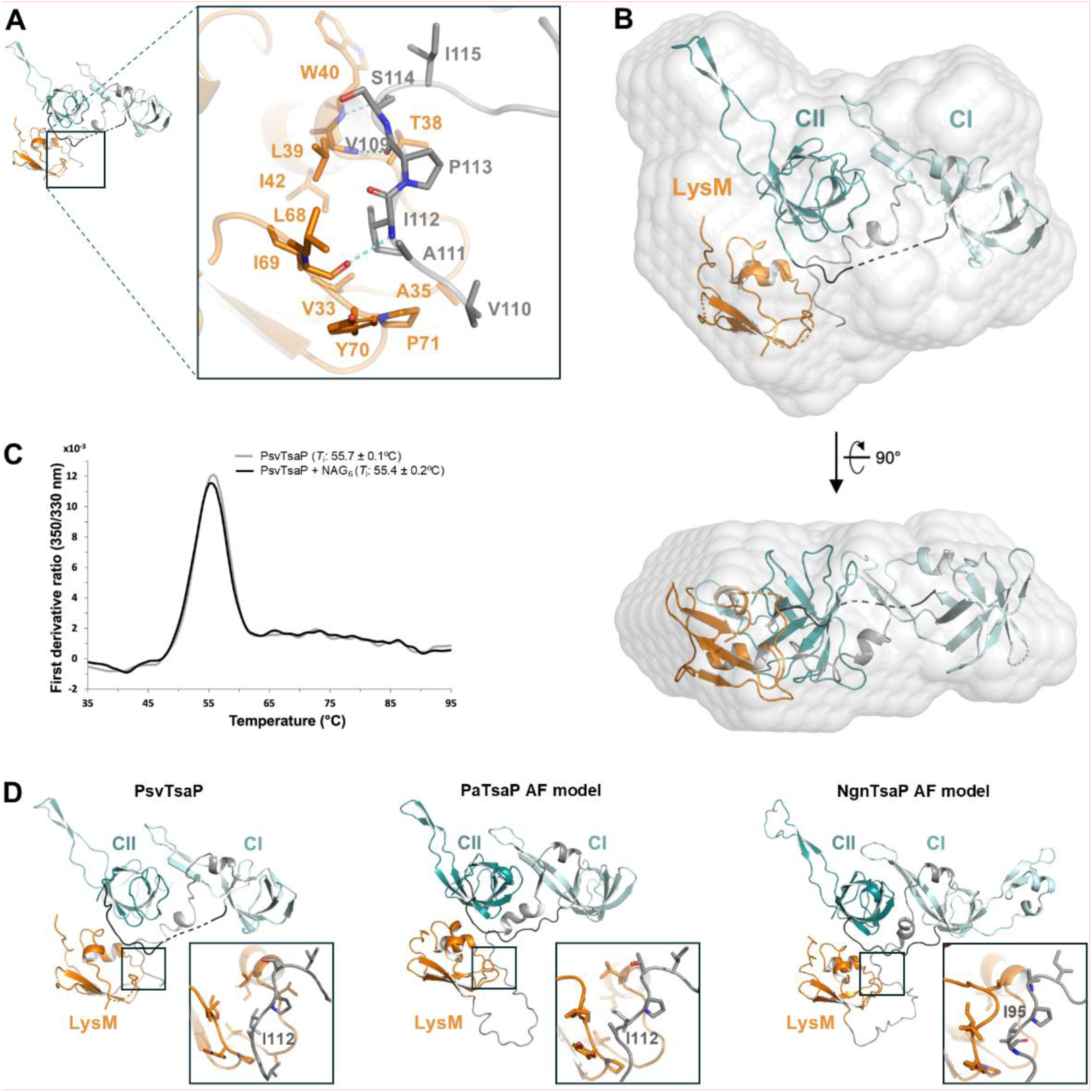
PG-locked architecture determined in PsvTsaP and in other TsaP homologues. (**A**) Close-up view of the PsvLysM:L1 inter-domain interface, showing a zoom around residues contributing to inter-domain contacts that are depicted as sticks. (**B**) Structural model of the mature full-length PsvTsaP determined in solution by SAXS. (**C**) NanoDSF first-derivative unfolding profiles of PsvTsaP in the absence and in presence of NAG_6_, and calculated *T*_i_ values. (**D**) Structural comparison PsvTsaP crystal structure with AF3 models of TsaP from *P. aeruginosa* (PaTsaP) and *N. gonorrhoeae* (NgnTsaP), showing zoom views of LysM:L1 contacts in each case. In all panels, protein domains are coloured as in Fig. 1.

To determine whether the PG-locked conformation observed in the crystal is maintained in solution, we performed SAXS analyses. The *ab initio* molecular envelope of PsvTsaP determined by SAXS exhibits an asymmetrical planar shape with approximate dimensions of 85×70×35 Å (Fig. 2B). The crystallographic model could be readily accommodated within the SAXS-derived envelope, and its calculated scattering profile showed good agreement with the experimental data (Fig. 2B and Appendix Fig. S2A, B). These data support the existence of a PG-locked conformation in solution, as observed in the crystallographic structure, compatible with the LysM:L1 interaction and hindrance of the PG-binding site. Next, we used nanoDSF to further explore the interaction of the full-length PsvTsaP with the PG analogue NAG_6_ in solution. These assays show identical *T*_i_ values for PsvTsaP in the absence (ligand-free control) and in the presence of NAG_6_ (Fig. 2C) indicating that, under the conditions tested, the ligand does not measurably alter the thermal stability of the full-length protein. Although our NAG_6_-binding experiments do not directly demonstrate binding to full-length PsvTsaP, they are consistent with a PG-locked conformation maintained both *in crystallo* and in solution. This suggest that binding of the PG analogue, and likely of PG itself, is insufficient to promote the conformational rearrangements required to expose the LysM domain for cell-wall engagement.

We also compared the PsvTsaP crystal structure with AlphaFold models of TsaP homologues PaTsaP and NgnTsaP (Fig. 2D). This comparison shows not only overall structure conservation of the PG-locked fold observed in PsvTsaP, but a similar LysM:L1 interaction blocking the PG-binding site of the PaLysM and NgnLysM domains (Fig. 2D). Collectively, our structural and biophysical data, together with comparative structural modelling, support a conserved PG-locked conformation in TsaP proteins. This conformation is compatible with restricted accessibility of the LysM-glycan binding site and supports a model in which LysM-mediated anchoring to the cell-wall is prevented until conformational associated with PilQ binding expose the PG-binding site.

### The LysM module of PsvTsaP displays conserved structural and biophysical features of a PG-binding domain

LysM domains are protein motifs that specifically recognize NAG-containing polysaccharides, such as bacterial PG or chitin present in the fungal cell-wall (Buist *et al*, 2008). They are widespread across bacterial, fungal and plant extracellular proteins, conferring the ability to interact with the bacterial and fungal cell surfaces required for their function (Buist *et al*, 2008). In the case of TsaP proteins, the LysM domain is presumed to bind PG to enable the T4P secretin anchoring to the bacterial cell wall. However, to our knowledge, verification of this interaction remains elusive (McCallum *et al*, 2021; Chang *et al*, 2016). Thus, we explored the interaction of PsvLysM with PG, starting with structural homology analysis to assess its degree of conservation (Fig. 3A, B). Structural superposition of PsvLysM with LysM domain structures determined in complex with oligosaccharides, including NlpC/P60 D,L-endopeptidase from *T. thermophilus* bound to NAG_6_ (TthNlpC/P60:NAG_6_; PDB: 4UZ3; (RMSD = 2.154 Å for 39 Cα atoms) (Wong *et al*, 2015) and LysM effector Ecp6 from *C. fulvum* bound to NAG_4_ (CflEcp6:NAG_4_; PDB: 4B8V; RMSD = 2.017 Å for 39 Cα atoms) (Sánchez-Vallet *et al*, 2013), indicates strong structural conservation of both the overall fold and the carbohydrate-binding site of PsvLysM (Fig. 3A). The latter includes a GDTL motif (aa 37-40 in PsvLysM) at the βA-αA loop, and a LIYPG motif (aa 68-72 in PsvLysM) at the αB-βB loop, which are involved in ligand interaction and are highly conserved across TsaP-LysM and other LysM domains inspected (Fig. 3A, B) (Buist *et al*, 2008). Based on these observations, we used nanoDSF experiments to explore the interaction of the PsvLysM domain with the NAG_6_ oligosaccharide, as PG analogue. Comparison of the *T*_i_ values measured for PsvLysM alone (51.5 ± 0.4°C) and in the presence of NAG_6_ (66.1 ± 0.9°C) showed a stabilization of the PsvLysM domain of ∼15°C in the presence of the oligosaccharide, supporting the interaction of PsvLysM with the PG analogue (Fig. 3C). Altogether, our homology and biophysical studies suggest that TsaP-LysM domains likely constitute a conserved PG-binding module consistent with the proposed role of TsaP in anchoring the T4P machinery to the bacterial cell wall. Moreover, our sequence and 3D-structure comparisons show that the PsvLysM and TsaP homologues inspected bear an additional C-terminal strand (βC in PsvLysM) (Fig. 3A, B), absent in typical LysM domains, thereby defining a distinctive LysM-domain architecture in T4P secretin-associated proteins.

**Figure 3.**
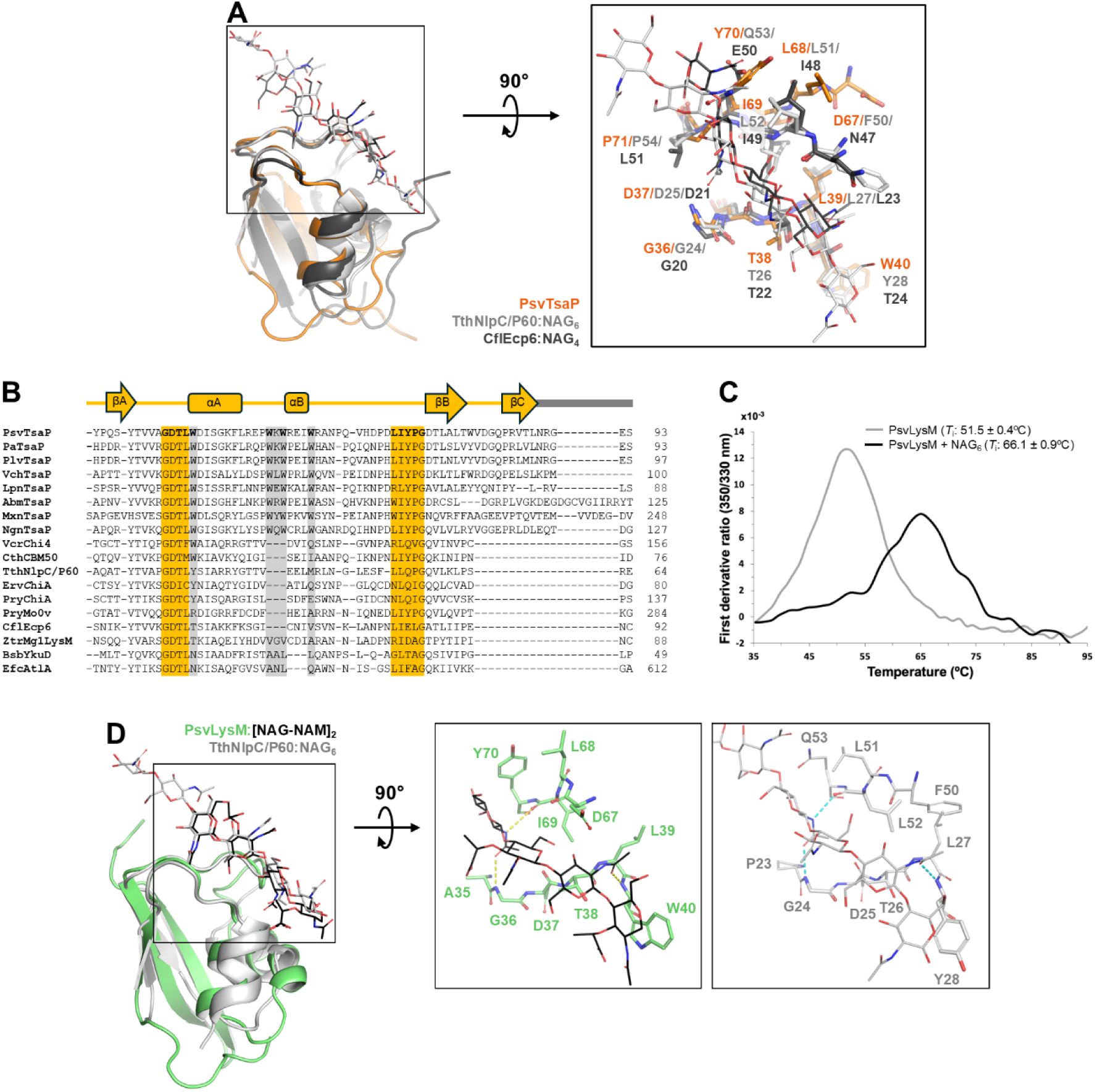
**Canonical and non-canonical features of the PsvLysM domain conserved in other TsaP-LysM domains**. (**A**) Structural superposition of PsvLysM (orange) with LysM domains of TthNlpC/P60:NAG_6_ (PDB: 4UZ3; in light grey) and CflEcp6:NAG_4_ (PDB: 4B8V; in dark grey). A zoom view of the PG-binding site is shown, with conserved residues involved in glycan interactions depicted as sticks. (**B**) Sequence alignment of PsvLysM and other LysM domains from TsaP homologues and representative LysM-containing proteins of known structure. Conserved residues at βA-αA and αB-βB ligand-binding motifs, including the GDTL and LIYPG regions, and in hydrophobic patches involved in inter-domain interfaces are highlighted in orange and grey, respectively. (**C**) NanoDSF first-derivative unfolding profiles corresponding to the isolated PsvLysM domain, in the absence and in presence of NAG_6_, and calculated *T*_i_ values. (**D**) AF3 structural model of the PsvLysM:[NAM-NAG]_2_ complex (in green and black) superimposed to TthNlpC/P60:NAG_6_ crystal complex. Insets show close-up views of the predicted glycan-binding region in PsvLysM and the PG-binding site in TthNlpC/P60. Conserved residues involved in ligand interactions and oligosaccharide molecules are shown as sticks.

To further explore the mode of interaction of PsvLysM with PG, we modelled the structure of PsvLysM bound to the PG analogue [NAM-NAG]_2_ using AF3. The computational complex, referred to as PsvLysM:[NAM-NAG]_2_, exhibits pTM (0.62) and ipTM (0.70) values that support the structural plausibility of the model. The analysis of this complex predicts that PsvLysM adopts a canonical LysM fold highly conserved with respect to the PsvTsaP crystal structure (Fig. 3D and Appendix Fig. S3A, B). The tetrasaccharide molecule is accommodated within the PG-binding site defined by the loops βA-αA and αB-βB. Superimposition of PsvLysM:[NAM-NAG]_2_ with TthNlpC/P60:NAG_6_ crystal complex (PDB: 4UZ3) predicts a conserved ligand-binding pose (Fig. 3D) and conserved of hydrogen-bonding interactions with the *N*-acetyl/hydroxyl groups of the oligosaccharide mediated by backbone atoms within the βA-αA and αB-βB loops (G36, W40 and I69 in PsvLysM; G24, Y28 and L52 in TthNlpC/P60) (Fig. 3D). Moreover, our PsvLysM:[NAM-NAG]_2_ complex predicts a closure of the αB-βB loop over the ligand, with respect to the PG-free PsvTsaP crystal structure. This rearrangement is consistent with a tighter LysM:ligand interaction, and it is also observed in experimental LysM:NAG_n_ complexes reported in the literature such as the TthNlpC/P60:NAG_6_ and CflEcp6:NAG_4_ complexes (Wong *et al*, 2015; Sánchez-Vallet *et al*, 2013) (Fig. 3A). Taken together, these observations support that PsvLysM conserves the PG recognition mode described for other LysM domains. However, despite the conservation observed, the PsvLysM:[NAM-NAG]_2_ complex also reveals a prominent aromatic patch (W40, W51, W53 and W57) adjacent to the PG-binding site that is highly and exclusively conserved in TsaP proteins (Fig. 3B). This patch contributes to the PsvTsaP interdomain interface, becoming partially (W53 and W57) and fully accessible (W40 and W51) to the solvent upon LysM release from the rest of the protein. Since exposed aromatic residues are typically involved in protein-protein and protein-ligand interactions, it is conceivable that this conserved hydrophobic patch may contribute to extend the canonical PG-binding site. Indeed, the PsvLysM:[NAM-NAG]_2_ model shows how the oligosaccharide interacts with W40 and extends toward W51. This observation highlights W51 as a potential extension of the carbohydrate binding site; a hypothesis that will require further studies to be corroborated. Altogether, our structural, biophysical, and modelling analyses support PsvLysM as a *bona fide* carbohydrate-binding LysM domain and suggest that PG recognition may involve an extended carbohydrate-binding site.

### PsvTsaP:PilQ interaction provides a structural basis for LysM-domain release and cell-wall engagement

Cryo-EM structures of PaT4P and MxnT4P show the TsaP-LysM domain positioned away from the protein core when bound to PilQ, in a conformation compatible with PG binding. In contrast, our data supports that PsvTsaP and TsaP reach the bacterial surface in a PG-locked conformation, raising the question of what triggers TsaP-LysM release. To address this, we modelled the PsvTsaP:PilQ complex using AF3. The interaction between TsaP and PilQ has been experimentally demonstrated in *P. aeruginosa*, where PaTsaP co-purifies with PilQ and a cryo-EM structure of the complex has been determined (PDB: 6VE2 (McCallum *et al*, 2021)). Our PsvTsaP:PilQ model converges with this experimental structure to an RMSD of 1.839 Å over 998 Cα atoms, indicating that AF3 accurately recapitulates the experimentally validated interaction geometry. This structural convergence, supported by pTM (0.70) and ipTM (0.73) confidence metrics, provides a robust framework for examining the molecular details of PsvTsaP:PilQ assembly (Appendix Fig. S4A, B, C).

Inspection of PsvTsaP:PilQ complex shows PsvTsaP to stack against β4, β7, β12 and β13 of PilQ β-barrel through β1-β2 and β8-β9 loops of CI, and β1’-β2’ and β8’-β9’ loops of CII, closely resembling the interactions mode observed in the PaTsaP:PilQ complex (Fig. 4A and Appendix Fig. S4C). Analysis of the PsvTsaP:PilQ interface indicates: i) two discrete complementary charged patches distributed across the interacting surfaces of PsvPilQ and PsvTsaP that would enable up to six salt-bridge interactions (Fig. 4B); and ii) three hydrophobic patches in PsvTsaP predicted to interact with three hydrophobic regions in PsvPilQ (Fig. 4C). Interestingly, most residues involved in these interactions exhibit high conservation across TsaP and PilQ homologues inspected (Appendix Fig. S1A and Appendix Fig. S4C), which supports the ability of PsvTsaP and other TsaP proteins to bind the secretin via a conserved pattern of electrostatic and hydrophobic interactions.

**Figure 4.**
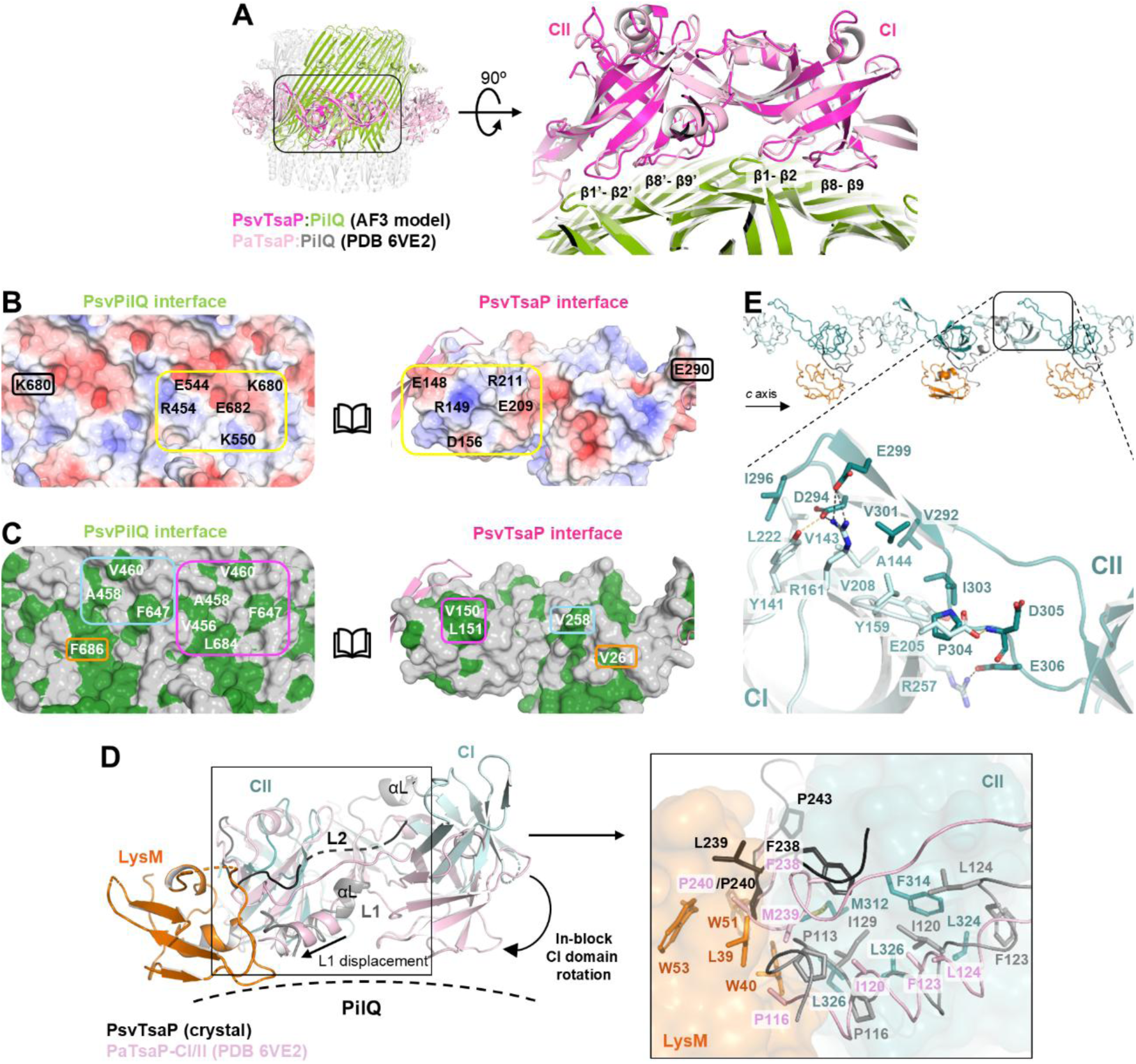
Structural remodelling of PsvTsaP triggered by PilQ interaction. (**A**) Structure of PsvLysM4:PilQ complex (in magenta and green) modelled with AF3, and superimposed to the PaTsaP:PilQ complex (PDB 6VE2; in pink and grey). The inset shows a close-up view of the interface formed by CI/II domains of LysM4/TsaP and PilQ, including labels for key LysM4/TsaP interacting loops: β1-β2, β1′-β2′, β8-β9 and β8′-β9′. (**B**) Open-book electrostatic potential surface representation of the modelled PsvTsaP:PilQ interface. Positively and negatively charged residues are coloured in blue and red, respectively, with those contributing to the complex interface highlighted with labels. (**C**) Open-book hydrophobic surface representation of the modelled PsvTsaP:PilQ interface. Hydrophobic residues contributing to TsaP:PilQ are indicated with labels. (**D**) Structural superimposition of PsvTsaP crystal structure onto PaTsaP cryo-EM structure (PDB: 6VE2) evidencing the in-block rotation of the CI domain and the displacement observed in L1. The right panel shows a close-up view of the PsvLysM:CI/CII interface, showing key residues involved in the structural rearrangement as sticks. PsvTsaP and PaTsaP molecules are coloured as in Fig. 1 and Fig. 4A, respectively. (**E**) Inter-monomer contacts along the *c* axis of the crystal packing representative of the interactions between CI and CII domains of PsvTsaP *in trans.* Residues involved in interdomain contacts are represented as sticks.

Structural comparison between the PilQ-free crystal structure of PsvTsaP and the PilQ-bound model allows us to propose the following activation mechanism. The comparison of the PaTsaP:PilQ complex with the PsvTsaP crystal structure, representing a PilQ-free state, shows a rotation of ∼35 degrees between CI and CII domains upon interaction of PsvTsaP with the secretin (Fig. 4D). This rearrangement is also predicted in our computational PsvTsaP:PilQ complex, and it enables PsvTsaP and PaTsaP to accommodate the curvature of the secretin channel (Appendix Fig. S4A). CI/CII-interdomain rotation is coupled to an in-block displacement of the L1 connector by >7 Å towards the LysM domain. This motion results, on the one hand, from a sliding of hydrophobic residues at the L1:CII interface (I115, I120, F123, L124 and I129 in PsvL1; and V264, V266, M312, F314, Y317, L320, Y322, L324, L326 in PsvCII) and, on the other hand, from the presence of αL and αĹ helices and three proline residues (P112, P116, and P140) framing the connector, which imposes a rigid architecture on L1 (Fig. 4D). CI/CII-interdomain rotation also propagates a displacement across the L2 connector by ∼2.5-4.5 Å. This movement primarily involves the shift of two phenylalanine residues (F227 and F238) at PsvTsaP interdomain interface, which pulls on the rigid proline-rich C-terminus of L2 fostering a destabilization of LysM:L2 interface. In this scenario, the rotation of CII forces L1 towards LysM while disrupting LysM:L2 interface, promoting the release of the LysM domain. Our analysis thus suggests that interaction of PsvTsaP with PilQ triggers a cascade of conformational changes leading to LysM release and exposure of its PG-binding site for cell-wall engagement. Thus, PsvTsaP:PilQ association may act not only as a structural anchor for PsvTsaP assembly, but also as a switch enabling PsvTsaP interaction with PG. This mechanism may be extended to other TsaP proteins based on the high degree of conservation observed in the hydrophobic contacts involved in the rearrangement of CI, L1 and L2 regions in PsvTsaP homologues inspected.

Finally, the PaT4P cryo-EM structure shows how PaTsaP:PilQ complex formation involves PaTsaP interactions *in trans* during its assembly onto the secretin. The latter consist of head-to-tail contacts between the CII β-hairpin of one monomer and the CI β-barrel core of the adjacent molecule, as *in cis* contacts between the CI and CII domains (McCallum *et al*, 2021). However, details of TsaP:TsaP inter-protomer interactions remain elusive due to the limited resolution of the PaT4P model. Notably, analysis of PsvTsaP crystal packing reveals inter-molecular contacts propagating along the *c* axis of the crystal lattice illustrating the head-to-tail interactions *in trans* observed for TsaP in the PaT4P structure (Fig. 4E and Appendix Fig. S4A). This interaction involves numerous hydrophobic residues forming a central core (Y141, V143, A144, Y159, V208, L222 in CI; and V292, I296, V301, I303, P304 in CII), which is surrounded by polar residues establishing hydrogen-bonding interactions (Y141, E205, D294, D305) and salt-bridge (R161, D294, R257, E306) contacts. Sequence analysis indicates strong conservation of the hydrophobic character of the residues forming the core of the interface, with most positions displaying only conservative substitutions (Y141, V143, A144, Y159, V208, I303, V292, I296 and P304). This aligns with a conserved mechanism of interaction *in trans* across PsvTsaP monomers, and other homologues examined, during TsaP assembly onto the secretin PilQ, which seems to be primarily guided by hydrophobic interactions.

## DISCUSSION

Our study shows that TsaP is not a constitutive PG tether but a regulated structural component whose cell-wall binding activity is controlled by its association with the T4P secretin PilQ. Previous work established that TsaP localizes around the PilQ secretin and contributes to the organization of the outer-membrane T4P subcomplex in both *M. xanthus* and *P. aeruginosa* (Siewering *et al*, 2014; Chang *et al*, 2016; McCallum *et al*, 2021). In these studies, the LysM domain projects separated from the CI/II core of TsaP, and toward the cell wall, to presumably interact with the bacterial PG. Unexpectedly, our structural characterization of full-length PsvTsaP, *in crystallo* and in solution, shows that this TsaP is not exported in a constitutively PG-binding-competent conformation. Instead, PsvTsaP adopts a compact arrangement in which the LysM domain is sequestered against the CI/CII core through L1-mediated intramolecular interactions, and we provide NanoDSF evidence consistent with the PG-locked conformation being unable to engage a PG analogue (Fig. 2A, C).

This observation has immediate mechanistic implications. The PG-binding site of the LysM domain is occluded by the L1 linker, which locks the domain in a non-productive arrangement also observed in the AlphaFold model of TsaP homologues from *P. aeruginosa* and *N. gonorrhoeae* (Fig. 2D). These results support the hypothesis that the PG-locked conformation reported represents a conserved resting state of TsaP-like proteins, and that TsaP is exported in an autoinhibited state that prevents indiscriminate engagement with the bacterial cell wall.

At the same time, our data indicates that the lack of glycan-binding by PsvTsaP is a result of this PG-conformation and not due to its LysM domain itself. Structural comparison with experimentally determined LysM:oligosaccharide complexes from TthNlpC/P60 and CflEcp6 shows that PsvLysM exhibits a canonical LysM fold and conservation of the PG-binding site, including the GDTL-containing βA-αA loop and the LIYPG-containing αB-βB loop (Buist *et al*, 2008; Sánchez-Vallet *et al*, 2013; Wong *et al*, 2015) (Fig. 3A, B). This is consistent with our computational PsvLysM:[NAM-NAG]_2_ complex modelled with AF3; and with nanoDSF assays showing the ability of the isolated PsvLysM domain to interact with the oligosaccharide NAG_6_, which confirms its function as a PG-binding module (Fig. 3C). These observations provide direct experimental support for a long-standing but untested assumption that TsaP-LysM domains contribute to PG recognition during T4P assembly (Siewering *et al*, 2014; Chang *et al*, 2016; McCallum *et al*, 2021). Our analysis of PsvLysM and other TsaP-associated LysM domains also show that, despite their general conservation, they exhibit distinct features with respect to previously characterized non-TsaP-LysM structures. This includes an extra C-terminal β-strand, and a prominent aromatic patch adjacent to the PG-binding site (Fig. 3B). The PsvLysM:[NAM-NAG]_2_ complex further supports the latter as a distinctive feature that may contribute to extending the PG-binding groove (Fig. 3D). Although this hypothesis requires experimental validation, it suggests that TsaP proteins have adapted a widespread carbohydrate-binding fold to the specific mechanical requirements of T4P anchoring through structural features not observed in other LysM motifs.

The central conceptual advance emerging from our study is a model in which PilQ binding acts as the structural trigger for TsaP activation. In our PsvTsaP:PilQ model, the CI/CII core engages the secretin through complementary electrostatic and hydrophobic patches that closely mirror those described for the PaPilQ:TsaP complex determined by cryo-EM (McCallum *et al*, 2021) (Fig. 4B, C). Comparison of this PilQ-bound conformation represented by the PsvTsaP:PilQ model and the cryo-EM structure of the PaPilQ:TsaP complex, with the PilQ-free crystal structure of PsvTsaP reveals a marked CI/CII rearrangement that propagates through the L1 and L2 connectors, leading to LysM-domain release and exposure of its PG-binding site (Fig. 4D). These observations have enabled us to propose a model in which the interaction with PilQ does not simply recruit TsaP to the outer-membrane complex but promotes exposure of the LysM domain, thereby licensing productive interaction with the cell wall. This provides a mechanistic explanation for how TsaP can couple secretin assembly to cell-envelope anchoring while avoiding ectopic PG binding in the free state.

Moreover, our crystal structure of PsvTsaP reveals head-to-tail intermolecular contacts that recapitulate the TsaP interactions *in trans* observed in PaTsaP:PilQ cryo-EM complex, in which the CII region of one monomer engages the CI region of the adjacent monomer (McCallum *et al*, 2021) (Fig. 4E). This has enabled access to higher resolution details of these interactions key for TsaP:PilQ complex assembly, and which are dominated by conserved hydrophobic contacts (Fig. 4C). These observations extend the mechanistic model beyond single-monomer activation and provide a plausible structural basis for how adjacent TsaP molecules interact during ring assembly around the secretin. More broadly, they support the idea that conformational activation, oligomerization and envelope anchoring are coordinated features of TsaP function rather than independent assembly events.

Several aspects of the model proposed here remain further validation. Likewise, the proposed contribution of the conserved aromatic patch to glycan recognition and extension of the PG-binding site of TsaP-LysM domains, as well as the functional relevance of specific residues involved in L1/L2 contacts in maintaining the PG-locked state, and in TsaP:TsaP interface, which will require targeted mutagenesis and functional reconstitution. These questions are also relevant from a mechanistic and applied perspective, as the conformationally regulated architecture of PsvTsaP may define structural surfaces that could, in principle, be targeted to interfere with its engagement within the PilQ-associated complex. In particular, the intramolecular LysM-CI/CII interface and the linker-mediated PG-locked state described here provide a framework for considering how perturbation of TsaP conformational control might compromise proper integration of the T4aP outer-membrane/periplasmic module. Information that might be relevant for the future development of bacteriostatics against *P. savastanoi* and other pathogenic Gram-negative bacteria.

In summary, our data support PsvTsaP functioning as a TsaP involved in T4P assembly, and propose a functional model in which PsvTsaP is remains in an autoinhibited PG-locked conformation, followed by activation upon PilQ interaction, leading to a remodelling of the CI/CII-linker architecture that triggers the release of the LysM domain for PG binding at the appropriate cellular location. Such a mechanism would couple secretin assembly, TsaP oligomerization and envelope anchoring into a single coordinated process (Fig. 5). We also provide evidence that TsaP-LysM motifs function as PG-binding domains, thereby experimentally supporting a long-standing assumption, while providing detailed view of TsaP interactions *in trans* and with the secretin, significantly expanding our understanding of T4P assembly, not only in Psv but also in other Gram-negative bacteria. These results provide a framework for understanding how bacteria can adapt small domains, such as LysM motifs, and integrate them into a complex nanomachines to fine-tune the architecture, spatial organization and structural dynamics of these assemblies.

**Figure 5.**
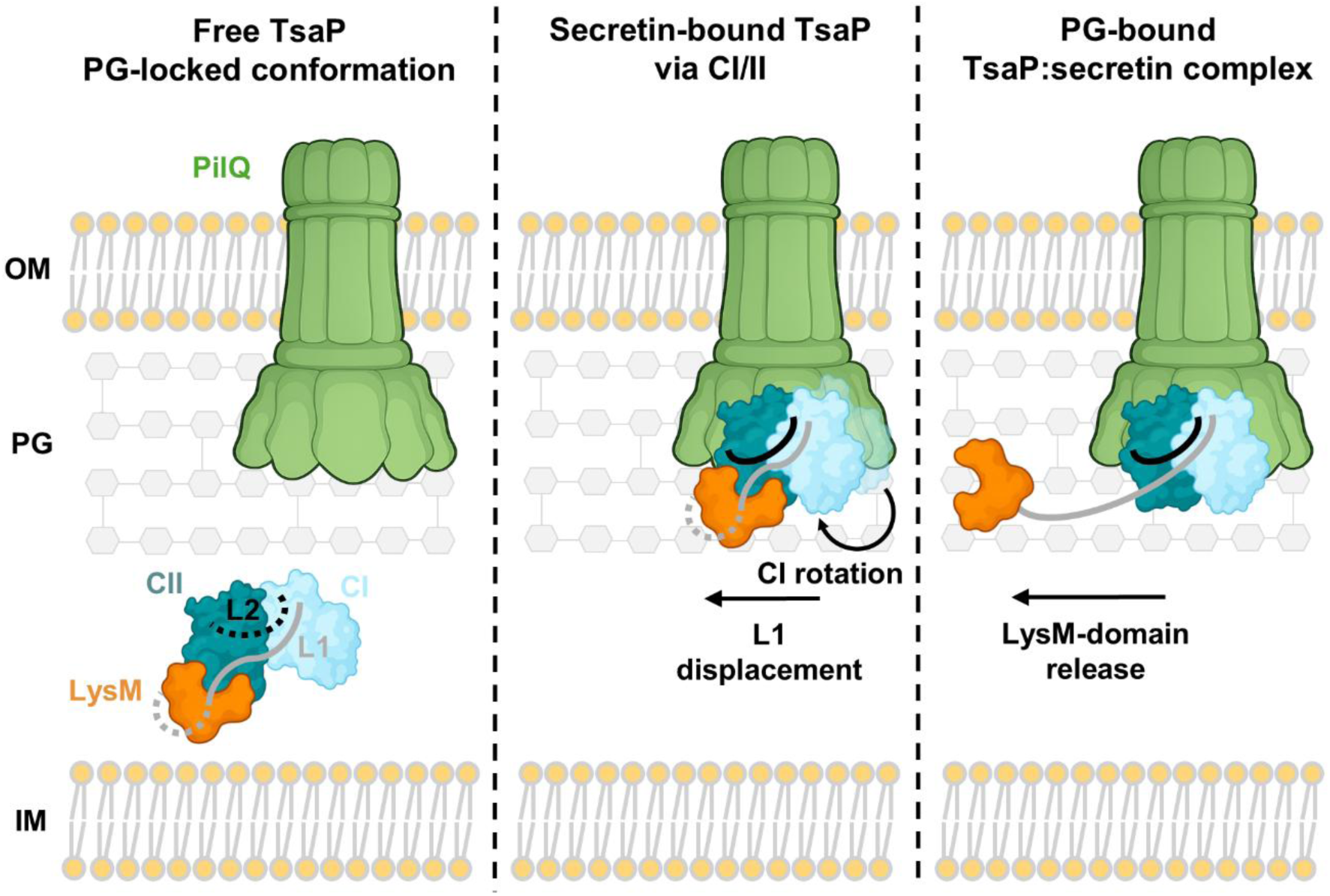
Model for conformational activation of PsvTsaP at the bacterial cell surface. The model poses PsvTsaP to be secreted in a PG-locked conformation, in which the LysM domain (in orange) stacks against the CII domain (in dark teal) via interactions of its PG-binding site and the L1 connector (in grey). The interaction with PilQ is proposed to induce rotation of the CI domain, followed by a major displacement of L1. This CI rotation triggers the detachment of the LysM domain, and the release of its PG-binding site, enabling the interaction of PsvLysM with the PG layer. OM, outer membrane; PG, peptidoglycan; IM, inner membrane.

## ACKNOWLEDGEMENTS

We thank Prof. Martín Martínez-Ripoll for maintaining the software infrastructure at the Department of Crystallography, Instituto de Química Física Blas Cabrera-Consejo Superior de Investigaciones Científicas (IQF-CSIC), which was essential for the X-ray crystallography studies presented in this work. We thank Prof. Bart Thomma for critically reading the manuscript.

## CONTRIBUTIONS

Major contributions to (i) the concept or design of the study (C.R., I.P-D., L. R-M.) (ii) the acquisition, analysis, or interpretation of the data (H.D-C., L.B-M., J.F-C., C.G-M., I.G.M., C.R., I.P-D., L.R-M.); (iii) writing of the manuscript (H.D-C., J.F-C., I.G.M., C.R., I.P-D., L.R-M.) and (iv) provision of resources (L.R-M., C.R., I.P-D.).

## DATA AVAILABILITY

The atomic coordinates and structure factors for the PsvTsaP(LysM4)/TsaP-like protein from *Pseudomonas savastanoi* pv. *savastanoi* NCPPB 3335 have been deposited in the Protein Data Bank (PDB) under accession code 31GV. All other data supporting the findings of this study are available within the Article and its Supplementary Information.

